# Chondroitin sulfate proteoglycan Windpipe modulates Hedgehog signaling in *Drosophila*

**DOI:** 10.1101/470096

**Authors:** Masahiko Takemura, Fredrik Noborn, Jonas Nilsson, Eriko Nakato, Tsu-Yi Su, Göran Larson, Hiroshi Nakato

## Abstract

Proteoglycans, a class of carbohydrate-modified proteins, often modulate growth factor signaling on the cell surface. However, the molecular mechanism by which proteoglycans regulate signal transduction is largely unknown. In this study, using a recently-developed glycoproteomic method, we found that Windpipe (Wdp) is a novel chondroitin sulfate proteoglycan (CSPG) in *Drosophila*. Wdp is a single-pass transmembrane protein with leucin-rich repeat (LRR) motifs and bears three CS sugar chain attachment sites in the extracellular domain. Here we show that Wdp modulates the Hedgehog (Hh) pathway. Overexpression of *wdp* inhibits Hh signaling in the wing disc, which is dependent on its CS chains and the LRR motifs. Conversely, loss of *wdp* leads to the upregulation of Hh signaling. Furthermore, knockdown of *wdp* increase the cell surface accumulation of Smoothened (Smo), suggesting that Wdp inhibits Hh signaling by regulating Smo stability. Our study demonstrates a novel role of CSPG in regulating Hh signaling.

## Introduction

Spatial and temporal regulation of growth factor signaling pathways is essential to proper development and disease prevention. Cell surface signaling events, such as ligand-receptor interactions, are often modulated by proteoglycans (D. Xu & Esko, 2014). Proteoglycans are carbohydrate-modified proteins that are found on the cell surface and in the extracellular matrix. They are composed of a core protein and one or more glycosaminoglycans (GAGs) covalently attached to specific serine residues on the core protein. GAGs are long, unbranched, and highly sulfated polysaccharide chains consisting of a repeating disaccharide unit. Based on the composition of the disaccharide units, proteoglycans are classified into several types, including heparan sulfate proteoglycans (HSPGs) and chondroitin sulfate proteoglycans (CSPGs).

HSPGs function as co-receptors by interacting with a wide variety of ligands and modulate signaling activities (Holt & Dickson, 2005; J.-S. Lee & Chien, 2004; Lindahl & Li, 2009; Poulain & Yost, 2015; D. Xu & Esko, 2014). *Drosophila* offers a powerful model system to study the functions of HSPGs *in vivo* because of its sophisticated molecular genetic tools and minimal genetic redundancy in genes encoding core proteins and HS synthesizing/modifying enzymes (Lander & Selleck, 2000; Nakato & Li, 2016; Perrimon & Bernfield, 2000; Takemura & Nakato, 2015). *In vivo* studies using the *Drosophila* model have shown that HSPGs orchestrate information from multiple ligands in a complex extracellular milieu and sculpt the signal response landscape in a tissue {Nakato & Li, 2016}. However, the molecular mechanisms of co-receptor activities of HSPGs still remain a fundamental question. Our previous studies predict that there are unidentified molecules involved in the molecular recognition events on the cell surface (Akiyama et al., 2008).

In addition to HS, *Drosophila* produces CS, another type of GAG (Toyoda, Kinoshita-Toyoda, & Selleck, 2000). CSPGs are well known as major structural components of the extracellular matrix. CSPGs have also been shown to modulate signaling pathways, including Hedgehog (Hh), Wnt, and fibroblast growth factor signaling (Cortes, Baria, & Schwartz, 2009; Townley & Bülow, 2018). Given the structural similarities between CS and HS, CSPGs may have modulatory, supportive and/or complementary functions to HSPGs. However, the mechanisms by which CSPGs function as a co-receptor are unknown. In contrast to a large number of studies on HSPGs, very few CSPGs have been identified and analyzed in *Drosophila* (Momota, Naito, Ninomiya, & Ohtsuka, 2011). Unlike HSPGs, CSPG core proteins are not well conserved between species (Olson, Bishop, Yates, Oegema, & Esko, 2006). Therefore, the identification of CSPGs cannot rely on the sequence homology to mammalian counterparts.

Recently, we have developed a glycoproteomic method to identify novel proteoglycans (Noborn et al., 2016; 2018; 2015). Briefly, this method includes trypsinization of protein samples, followed by enrichment of glycopeptides using strong anion exchange (SAX) chromatography. After enzymatic digestion of HS/CS chains, the glycopeptides bearing a linkage glycan structure common to HS and CS chains are identified using nano-liquid chromatography-tandem mass spectrometry (nLC-MS/MS). This method has successfully identified novel CSPGs in humans (Noborn et al., 2015) and *Caenorhabditis elegans* (Noborn et al., 2018).

To study the function of CSPGs in signaling, we applied the glycoproteomic method to identify previously unrecognized CSPGs in *Drosophila*. We found that Windpipe (Wdp) is a novel CSPG and affects Hh signaling. Overexpression of *wdp* inhibits Hh signaling in the wing disc. This inhibitory effect of Wdp on Hh signaling is dependent on its CS chains and LRR motifs. Consistent with the overexpression analysis, loss of *wdp* increases Hh signaling. Loss of *wdp* also increases cell surface accumulation of Smoothened (Smo), the Hh signaling transducer. Therefore, we propose that Wdp downregulates Hh signaling by disrupting cell surface accumulation of Smo.

## Results

### A glycoproteomic approach identified Wdp as a novel *Drosophila* CSPG

We investigated the potential presence of CSPGs in *Drosophila* using our recently-developed glycoproteomic approach that identifies core proteins and its CS attachment sites. A general workflow for the sample preparation, CS-glycopeptide enrichment, LC-MS/MS analysis and the subsequent data analysis is shown in Fig. 1A. Brifely, *Drosophila* third-instar larvae were collected from two different genotypes (wild type [Oregon-R] and a loss-of-function mutant for *tout-velu* [*ttv^524^*]) and the material was homogenized in ice-cold acetone. *ttv* encodes a *Drosophila* HS polymerase, and *ttv* mutants lack HS chains (Toyoda et al., 2000). The samples were incubated with trypsin and then passed over an anion exchange column equilibrated with a low-salt buffer. This procedure enriches for CS-attached glycopeptides as the matrix retains anionic polysaccharides and their attached peptides, whereas neutral or positively charged peptides flow through the column. The bound structures were eluted stepwise with three buffers of increasing sodium chloride concentrations. The resulting fractions were treated with chondroitinase ABC. This procedure reduces the lengths of the CS chains and generates a residual hexasaccharide structure still attached to the core protein. The chondroitinase-treated fractions were analyzed with positive mode nLC-MS/MS and an automated search strategy was used to identify CS modified peptides in the generated data set (Noborn et al., 2015).

**Figure 1.**
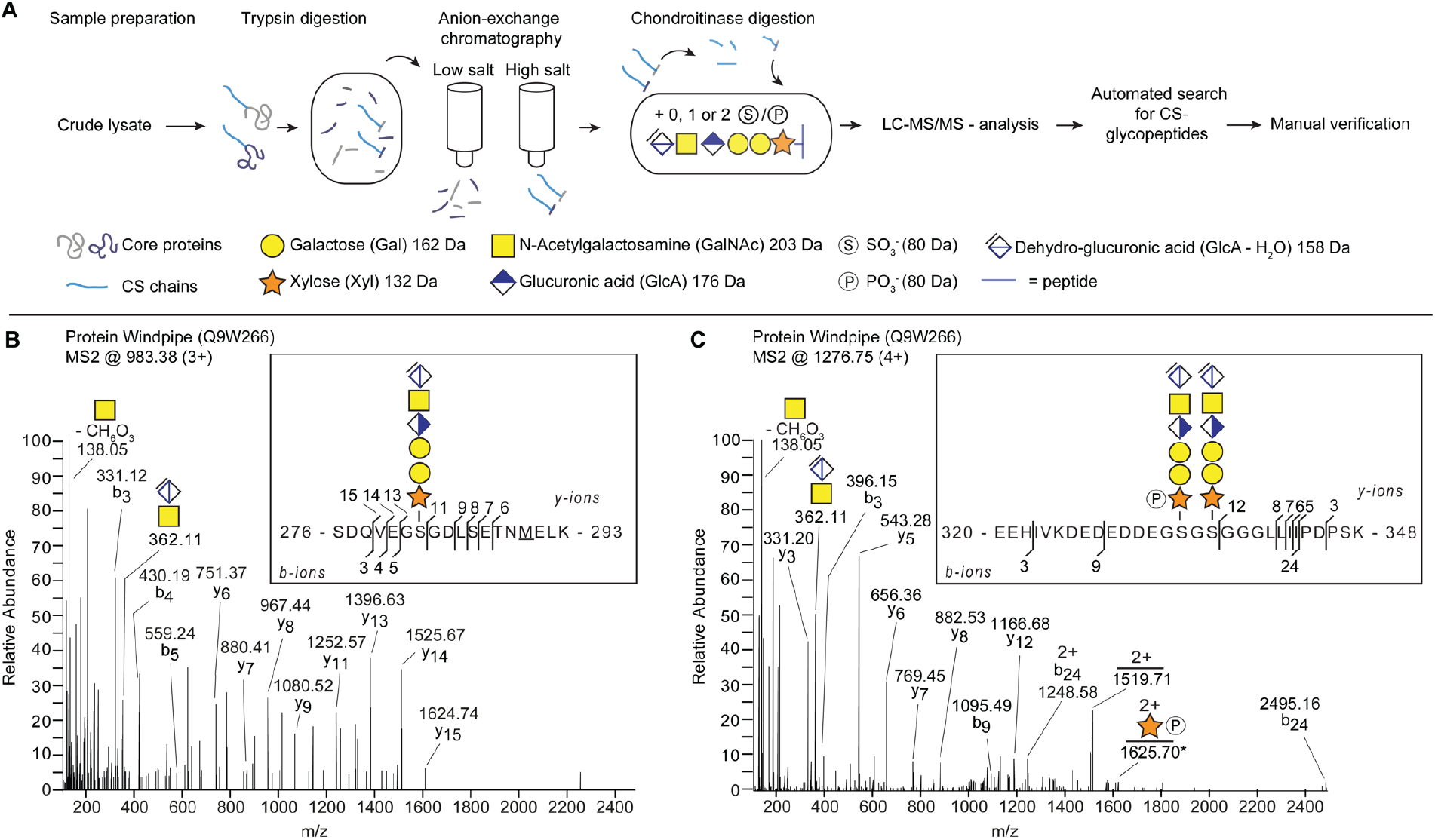
Identification of Wdp as a novel CSPG in *Drosophila*. (**A**) A scheme for identifying CSPGs in *Drosophila*. The workflow includes the enrichment of proteoglycans from fly extract, enzymatic hydrolysis and subsequent analysis and interpretation of mass spectra. (**B** and **C**) MS2 fragment mass spectra of Wdp protein (UniProt: Q9W266) showing two unique CS-glycopeptides. (B) Peptide (SDQVEGSGDLSETNMELK) identified with one hexasaccharide structure and one methionine oxidation (*m/z* 983.38; 3+) (C) Peptide (EEHIVKDEDEDDEGSGSGGGLLIIPDPSK) identified with two hexasaccharide structures where one of the hexasaccharides were modified with one phosphate modification (*m/z* 1276.76; 4+). The asterisk denotes the second isotopic peak.

The analysis revealed the Windpipe (Wdp) protein as a novel CSPG, which was modified with three CS-polysaccharides on two unique peptides (Fig. 1B and 1C). We detected Wdp glycopeptides from both wild-type and *ttv* mutant samples, further supporting that Wdp bears CS chains, not HS. One of the identified precursor ions (*m/z* 983.38; 3+) equated to the mass of a peptide with a SDQVEGSGDLSETNMELK sequence, derived from the middle part of the protein (amino acids 276– 293) (Fig. 1B). The peptide was modified with one hexasaccharide structure and one methionine oxidation. The measured mass (2947.1186 Da) deviated - 3.27 ppm from the theoretical value. The other identified precursor ion (*m/z* 1276.76; 4+) equated to the mass of a peptide with a EEHIVKDEDEDDEGSGSGGGLLIIPDPSK sequence, located in proximity to the previous peptide (amino acids 320–348) (Fig. 1C). The peptide was found to be modified with two hexasaccharide structures and where one of the hexasaccharides were modified with one phosphate modification. The measured mass (5102.9389 Da) deviated +3.05 ppm from the theoretical value. Detailed inspection of the spectra revealed several b- and y-ions as well as the prominent diagnostic oxonium ion at *m*/*z* 362.1, corresponding to the disaccharide [GlcAGalNAc-H_2_O+H]^+^ (Fig. 1B and 1C). Furthermore, one of the glycans in Fig 1C was found modified with one phosphate group at a xylose residue (peptide + xylose + phosphate, *m/z* 1625.70; 2+).

Wdp is a single-pass transmembrane protein containing four leucine rich repeat (LRR) motifs in the extracellular domain (Huff, Kingsley, Miller, & Hoshizaki, 2002). The three CS attachment sites (S282, S334, and S336) revealed by our glycoproteomic analysis are located in the extracellular domain (Fig. 3A). Interestingly, a recent study reported that Wdp negatively regulates JAK–STAT signaling by promoting internalization and lysosomal degradation of the Domeless (Dome) receptor (W. Ren et al., 2015). We further investigated the role of Wdp, a novel CSPG, in signal transduction.

### Overexpression of *wdp* inhibits Hh signaling

The growth and patterning of the *Drosophila* wing are controlled by multiple signaling pathways, including Decapentaplegic (Dpp; the *Drosophila* BMP), Wingless (Wg; the *Drosophila* Wnt), and Hedgehog (Hh) signaling (Baena-Lopez, Nojima, & Vincent, 2012; Tabata & Takei, 2004). To determine the role of *wdp* in these developmental signaling pathways, we first asked whether overexpression of *wdp* affects adult wing morphology. When *wdp* was overexpressed in the wing pouch using *Bx^MS1096^-GAL4* (Capdevila & Guerrero, 1994) (*Bx^MS1096^>wdp*), the wing size was reduced compared to that of control flies (*Bx^MS1096^>*) (Fig. 3C; compared to Fig. 3B). In addition, the distance between longitudinal wing veins 3 and 4 (L3 and L4) was aberrantly narrower. This decreased distance between L3 and L4 is indicative of reduced Hh signaling during wing development (Mullor, Calleja, Capdevila, & Guerrero, 1997; Strigini & Cohen, 1997).

Hh is produced in the posterior compartment of the wing disc and spreads towards the anterior compartment where Hh signaling induces target genes expression in a concentration-dependent manner (Briscoe & Thérond, 2013; Gradilla & Guerrero, 2013; Hartl & Scott, 2014). Expression of high-threshold target genes, such as Patched (Ptc; the Hh receptor) (Capdevila, Pariente, Sampedro, Alonso, & Guerrero, 1994) and Engrailed (En) (Patel et al., 1989) are induced in anterior cells near the anteroposterior compartment boundary by high levels of Hh signaling (Jia, Tong, Wang, Luo, & Jiang, 2004) (Fig. 2A and 2E). Low levels of Hh signaling induce the expression of *dpp* and the accumulation of full-length Cubitus interruptus (Ci; the transcriptional factor of Hh signaling) in a broader region (more distant away from the anteroposterior boundary) (Fig. 2A and 2C). To determine if Hh signaling is indeed affected by *wdp*, we overexpressed *wdp* in the dorsal compartment of the wing disc using *ap-GAL4* (Calleja, Moreno, Pelaz, & Morata, 1996; O’Keefe, Thor, & Thomas, 1998). We found that *wdp* overexpression in the dorsal compartment reduced the expression domains of both “high-threshold” targets (Ptc and En) and “low-threshold” targets (*dpp-lacZ^10638^*, a reporter for *dpp* expression, and full-length Ci) compared to those in the ventral compartment (Fig. 2B, 2D, and 2F). Notably, overexpression of *wdp* did not affect the pattern of a *hh* transcriptional reporter *hh-lacZ^P30^* (J. J. Lee, Kessler, Parks, & Beachy, 1992) (Fig. 2F). Together, *wdp* acts as a negative regulator of Hh signaling without affecting *hh* transcription

**Figure 2.**
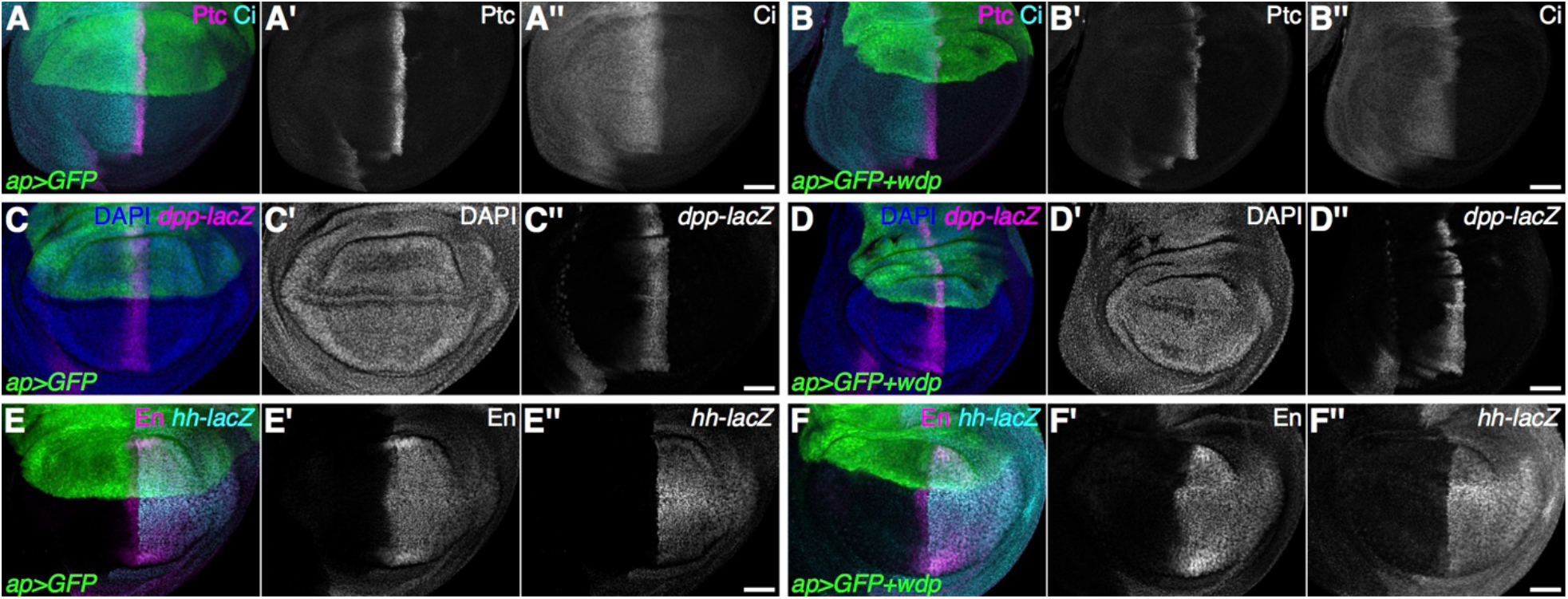
Overexpression of *wdp* reduces the Hh-signaling-active domain. (**A**–**F**) Control wing discs (A, C, and E) and wing discs overexpressing *wdp* with *ap-GAL4* (*ap>GFP+wdp*) (B, D, and F) were immunostained for the expression of Ptc, Ci (A and B), *dpp-lacZ* (C and D), En, and *hh-lacZ* (E and F). The expression domains of Ptc, Ci, and *dpp-lacZ* were reduced by *wdp* overexpression in the dorsal compartment compared to those in the ventral compartment. En expression induced by high-level Hh signaling in the anterior compartment is diminished by *wdp* overexpression. Note that the *hh-lacZ* expression is not affected by *wdp* overexpression. Nuclei were stained with DAPI (C and D). Anterior to the left; dorsal to the top. Scale bars: 50 µm.

On the other hand, Wdp does not appear to affect Dpp and Wg pathways. When *wdp* is overexpressed using *ap-GAL4* or *hh-GAL4* (a posterior compartment-specific GAL4 driver) (Tanimoto, Itoh, Dijke, & Tabata, 2000), we did not observe apparent defects in Dpp signaling activity, which was monitored by the expression of phosphorylated Mad (pMad) and Spalt major (Salm) (readouts of Dpp signaling). Similarly, no changes in expression of Senseless (Sens) and Distal-less (Dll) (readouts of Wg signaling) were detected (Fig. S1). These results are consistent with a previous report (W. Ren et al., 2015).

We also found that overexpression of *wdp* induces massive apoptosis, as detected with anti-cleaved Caspase-3 antibody (Fig. S2B). This likely contributed to the smaller adult wing phenotype observed in *Bx^MS1096^>wdp* flies. It was recently reported that Hh signaling is required for cell survival in wing disc cells (Lu, Wang, & Shen, 2017). To determine whether reduced Hh signaling is responsible for the observed apoptosis, we first asked if reduced Hh signaling results in apoptosis. We inhibited Hh signaling either by expressing an RNAi construct targeting *smo* (TRiP.HMC03577) (Fig. S2E), or by overexpressing *ptc* in the dorsal compartment using *ap-GAL4*. We found that neither treatment caused massive apoptosis (Fig. S2F and S2G), indicating that reduced Hh signaling is not sufficient to induce massive apoptosis in the wing disc. Furthermore, coexpression of a constitutively active form of Smo with Wdp did not suppress apoptosis in the wing disc (Fig. S2H). Thus, these results suggest that overexpression of *wdp* induces apoptosis, independent of reduced Hh signaling.

### CS and LRR motifs are necessary for Wdp to inhibit Hh signaling

Next, we asked whether the CS chains of Wdp are required for its function. In a CSPG core-protein, CS is attached to specific serine residues in the consensus serine-glycine dipeptide surrounded by acidic amino acids (Esko & Zhang, 1996). We generated a *UAS-wdp^ΔGAG^* construct in which all three serine residues (S282, S334, and S336) are substituted with alanine residues so that CS cannot be attached to the core protein (Fig. 3A). The *UAS-wdp^ΔGAG^* construct was inserted in the same genomic location (ZH-86Fb; (Bischof, Maeda, Hediger, Karch, & Basler, 2007)) as *UAS-wdp* using the *phiC31* site-specific integration system (Groth, Fish, Nusse, & Calos, 2004) in order to ensure the same expression level of the UAS transgenes.

**Figure 3.**
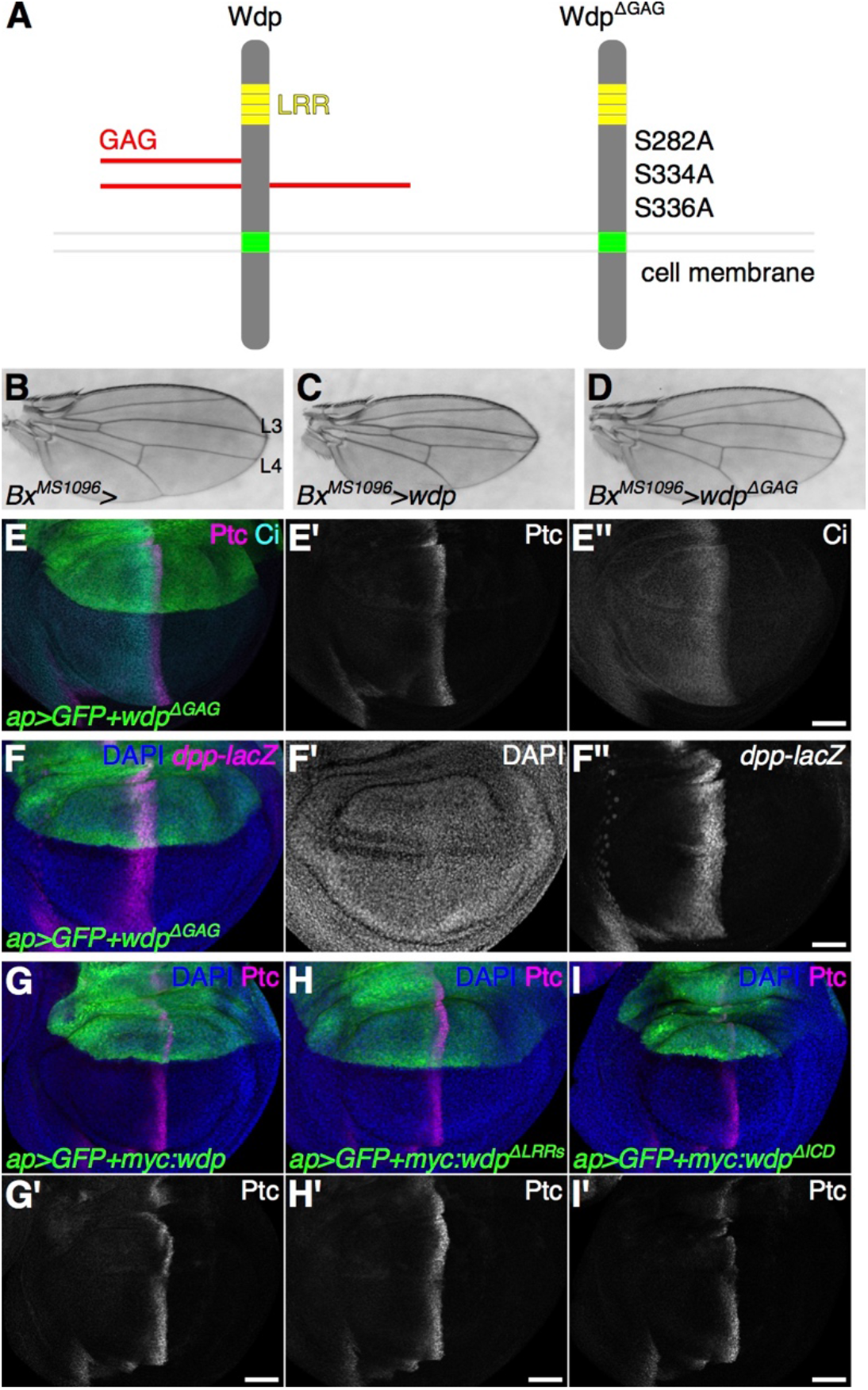
Wdp negatively regulates Hh signaling in a GAG-dependent manner. (**A**) A schematic drawing of wild-type Wdp and a mutant form of Wdp (Wdp^ΔGAG^). (**B**–**D**) A control adult wing (B) and adult wings expressing *UAS-wdp* (C) or *UAS-wdp^ΔGAG^* (D) with *Bx^MS1096^-GAL4*. (**E** and **F**) Wing discs expressing *UAS-wdp^ΔGAG^* with *ap-GAL4* were immunostained for Ptc, Ci (E) and *dpp-lacZ* (F). (**G**–**I**) Wing discs expressing *UAS-3xMyc:wdp* (G), *UAS-3xMyc:wdp^ΔLRRs^* (H), and *UAS-3xMyc:wdp^ΔICD^* (I) were immunostained for Ptc. Nuclei were stained with DAPI. Scale bars: 50 µm.

We found that *Bx^MS1096^>wdp^ΔGAG^* adult wings did not display the reduction in the distance between L3 and L4 (Fig. 3B). Consistent with this, the expression of Ptc, En, Ci, and *dpp-lacZ* in the wing disc were not affected by *wdp^ΔGAG^* overexpression in the dorsal compartment of the wing disc (Fig. 3E–G). These results indicate that CS chains are required for Wdp’s activity to downregulate Hh signaling.

To determine whether the LRR motifs and/or the intracellular domain of Wdp are necessary for inhibiting Hh signaling, we generated several Myc-tagged mutant constructs (Fig. S3) and examined their activities. Consistent with the earlier result (Fig. 2B), expression of a Myc-tagged Wdp (Myc:Wdp) led to the narrower Ptc expression domain (Fig. 3G). A mutant *wdp* construct lacking LRR motifs (Myc:Wdp^ΔLRRs^) failed to inhibit Hh signaling (Fig. 3H). On the other hand, a truncated construct lacking the intracellular domain (Myc:Wdp^ΔICD^) retained the ability to inhibit Hh signaling (Fig. 3I). Thus, in addition to CS chains, the LRR motifs of Wdp are required for inhibiting Hh signaling.

### Wdp expression in the wing disc

To monitor Wdp expression, we generated transgenic flies (*wdp^KI.HA^* and *wdp^KI.OLLAS^*) expressing epitope-tagged Wdp protein from its endogenous locus. We inserted a spaghetti monster GFP with 10 copies of HA or OLLAS tags (Nern, Pfeiffer, & Rubin, 2015; Viswanathan et al., 2015) near the C-terminus of Wdp (after Q652; Fig. 4A) using CRISPR–Cas9-mediated homology-directed repair (Gratz et al., 2014; X. Ren et al., 2014). The Wdp:HA expression was detected in the eye disc, adult midgut, and tracheal system (Fig. S3), consistent with previous reports (Huff et al., 2002; W. Ren et al., 2015).

**Figure 4.**
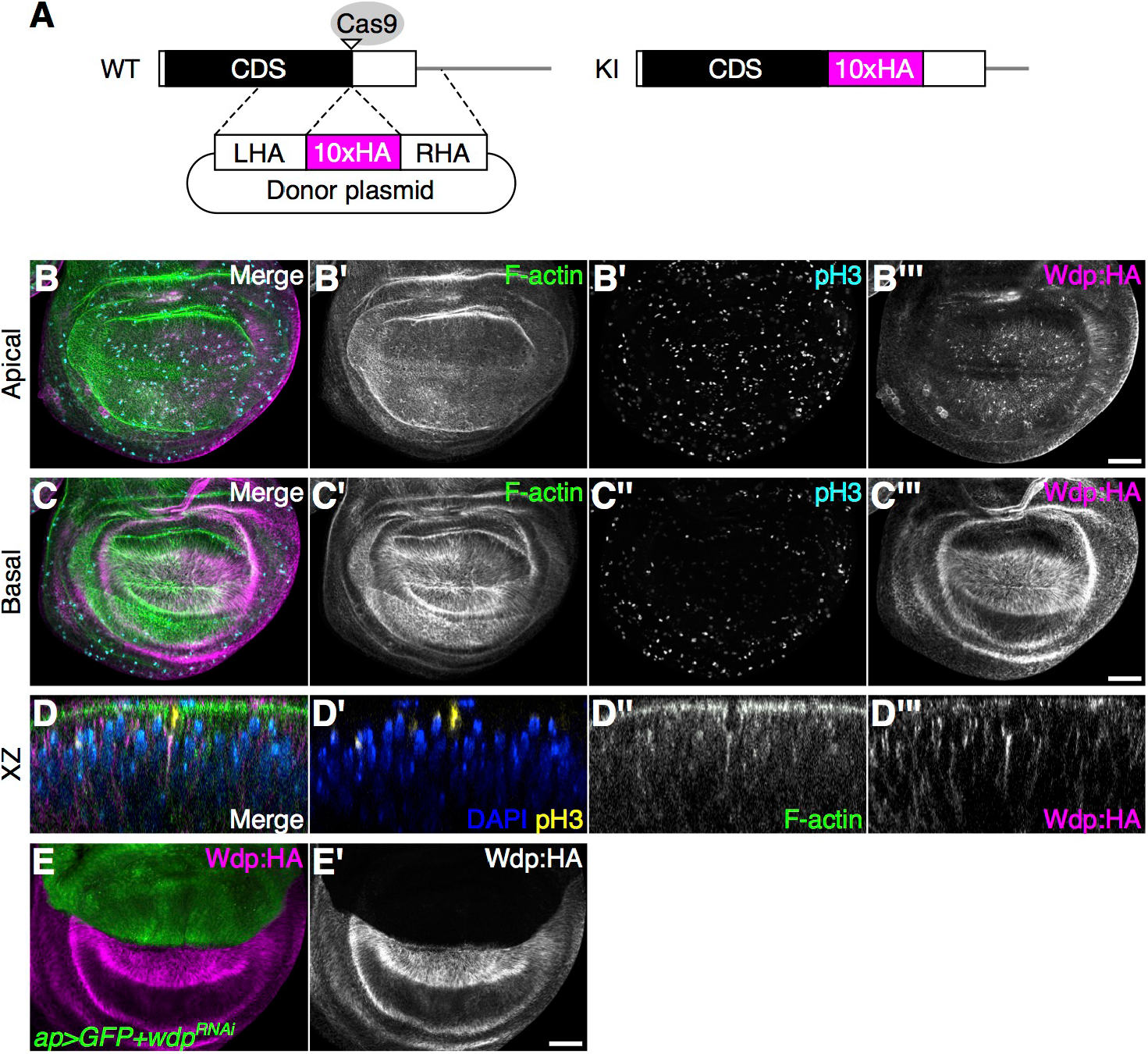
Wdp expression in the wing disc. (**A**) A schematic of CRISPR–Cas9-mediated gene editing of *wdp* for generating *wdp^KI.HA^*. Ten copies of an HA epitope tag (smGFP-HA) were inserted in frame near the stop codon of the *wdp* coding sequence (CDS). Only the last exon is shown. CDS, the black box; smGFP-HA, the magenta box; LHA, left homology arm; RHA, right homology arm; Cas9 target site, the open triangle. (**B**–**D**) Wing discs homozygous for *wdp^KI.HA^* were stained with Alexa Fluor 568-conjugated phalloidin (F-actin), anti-phospho histone H3 antibody (pH3, mitotic nuclei), and anti-HA antibody. Apical (B) and basal (C) sections of the same disc are shown. Intense staining of Wdp:HA was observed on the basal side of wing disc epithelium (C–C’’’). An optical cross section shows the accumulation of Wdp:HA in the basal projection of apically translocating mitotic cells (D–D’’’). (**E**) Wdp:HA is not detectable in the dorsal compartment of a wing disc expressing *wdp^RNAi^* (TRiP.HMC06302) with *ap-GAL4*. Nuclei were stained with DAPI (D and E). Scale bars: 50 µm.

In the wing disc, Wdp:HA is expressed in most of the wing disc cells with enrichment in the basal side, as detected by anti-HA antibody (Fig. 4B and 4C). This result was confirmed by anti-OLLAS antibody staining of the *wdp^KI.OLLAS^* wing discs (Fig. S3A and S3B). In the wing disc epithelium, mitotic nuclei apically translocate, but the cells maintain contact with the basal lamina via actin-rich basal projection (Ragkousi & Gibson, 2014). Interestingly, Wdp:HA is strongly enriched in such basal projections (Fig. 4A and 4D). However, physiological significance of this localization of Wdp in the basal projections of mitotic cells is unknown.

### Loss of *wdp* leads to higher levels of Hh signaling

To determine whether loss of *wdp* affects Hh signaling activity, we examined the effect of *wdp* RNAi knockdown in the wing disc. Expression of a *wdp^RNAi^* construct (TRiP.HMC06302) using *ap-GAL4* in *wdp^KI.HA/+^* flies led to the loss of Wdp:HA staining specifically in the dorsal compartment (Fig. 4E), validating the efficacy of RNAi-mediated knockdown of *wdp*. We then examined the effect of *wdp* knockdown on Hh signaling using the Ptc expression level as a readout of the Hh signaling activity. In control wing discs (*ap>FLP*), the signal intensity of Ptc staining in the dorsal compartment is comparable to that in the ventral compartment (Fig. 5B). On the other hand, *wdp^RNAi^* expression using *ap-GAL4* increased the signal intensity of Ptc staining only in the dorsal compartment (Fig. 5A and 5C). In addition, we observed that the *dpp-lacZ* expression domain was expanded anteriorly by *wdp* knockdown (Fig. 5D). In the adult wing, knockdown of *wdp* slightly expanded the distance between wing vein L3 and L4 near the distal tip (Fig. 5J, compared to Fig. 5I). Thus, *wdp* RNAi knockdown results in a moderate increase in Hh signaling.

**Figure 5.**
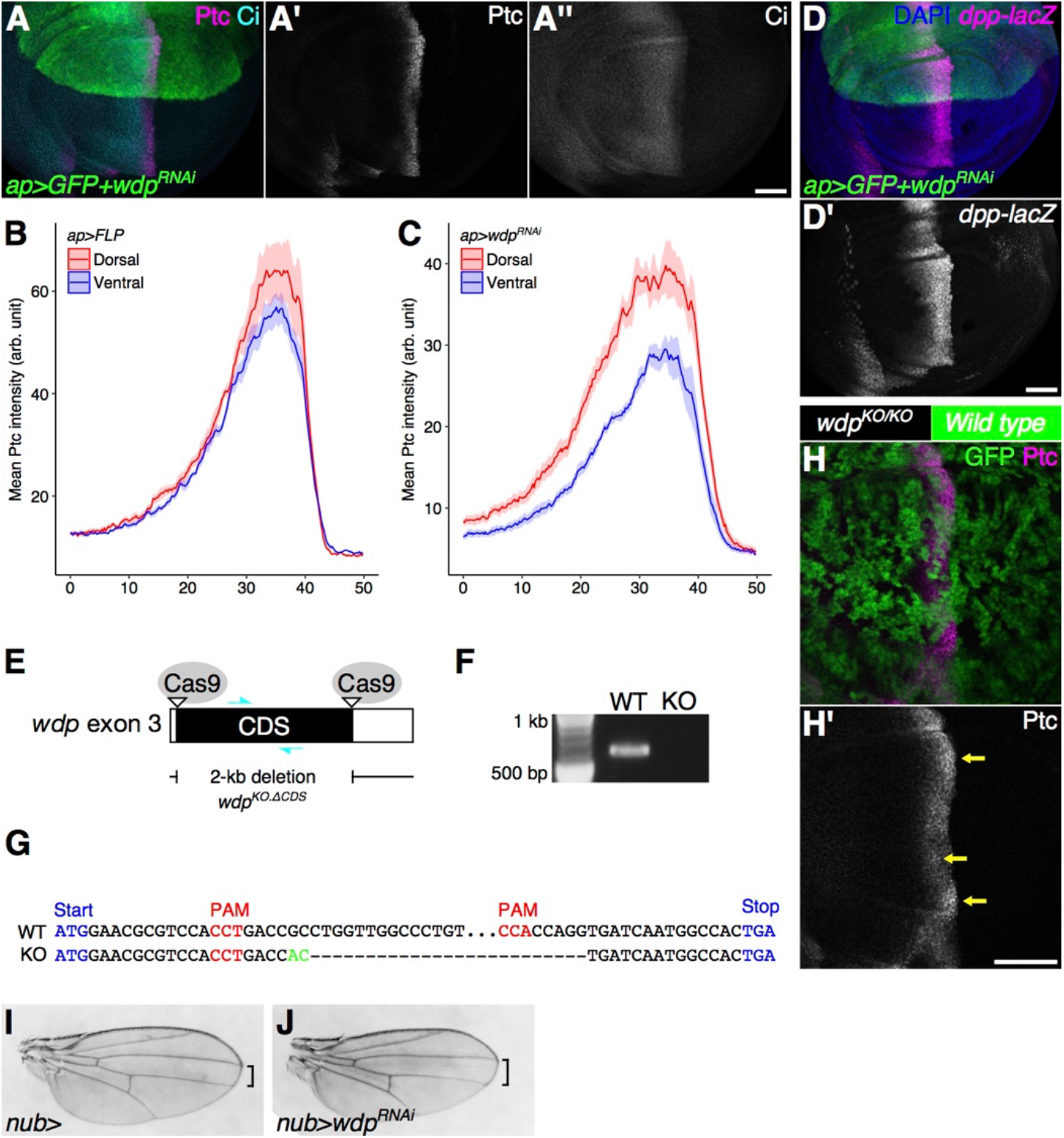
Loss of *wdp* leads to increased Hh signaling. (**A**) A wing disc expressing *UAS-wdp^RNAi^* (TRiP.HMC06302) with *ap-GAL4* was immunostained for Ptc and Ci. (**B** and **C**) Signal intensity plots of the Ptc expression in the dorsal compartment (red) and ventral compartment (blue) in wing discs expressing *UAS-FLP* (B) or *UAS-wdp^RNAi^* (C). Solid lines indicate the average intensity of Ptc staining and shaded areas show the standard error of the mean. (**D**) A wing disc expressing *UAS-wdp^RNAi^* with *ap-GAL4* was immunostained for *dpp-lacZ*. Nuclei were stained with DAPI. (**E**) A schematic of the generation of a *wdp* loss-of-function allele (*wdp^KO.ΔCDS^*) lacking most of the *wdp* CDS using the CRISPR– Cas9 system. (**F**) A PCR-based genotyping result for the wild-type (WT) and *wdp^KO.ΔCDS^* allele (KO) using a primer set shown as cyan arrows in E. (**G**) Genomic sequence of the *wdp* endogenous locus targeted by CRISPR–Cas9 to delete most of the *wdp* CDS. A small insertion is shown in green. (**H**) Somatic mosaic clones of *wdp^KO^* were induced in the wing pouch using *nub-GAL4 UAS-FLP*. Homozygous *wdp^KO^* mutant cells are marked by loss of GFP. Increased Ptc expression was observed in *wdp* mutant clones (yellow arrows) (**I**, **J**) A control adult wing (I) and a wing expressing *UAS-wdpRNAi* (TRiP.HM05118) (J) by *nub-GAL4*. Scale bars: 50 µm.

To confirm the *wdp* knockdown phenotypes, we generated a loss-of-function allele of *wdp* (*wdp^KO.ΔCDS^*), in which most of the *wdp* coding sequence was removed using CRISPR–Cas9-mediated defined deletion (Gratz et al., 2013) (Fig. 5E–G). *wdp^KO.ΔCDS^* homozygous mutant clones were induced in the wing pouch using the FLP–FRT system with *nubbin (nub)-GAL4 UAS-FLP* and their effect on Hh signaling was examined using anti-Ptc antibody. Consistent with the RNAi knockdown results, we observed a modest increase of Ptc expression in cells mutant for *wdp* (Fig. 5H). Taken together, we conclude that *wdp* negatively regulates Hh signaling in the *Drosophila* wing.

### Wdp inhibits Smo cell surface accumulation

The seven-pass transmembrane protein Smo is a key transducer of Hh signaling. In the absence of Hh, Ptc inhibits the phosphorylation of Smo, which is internalized and degraded (Zhu, Zheng, Suyama, & Scott, 2003). In the presence of Hh, restriction of Ptc on Smo is relieved, allowing Smo to accumulate on the cell surface and activate Hh signaling. Although *smo* transcription is ubiquitous, Smo protein expression levels are high in the posterior compartment of the wing disc where Ptc is not expressed (Fig. 6A) (Denef, Neubüser, Perez, & Cohen, 2000). We found that knockdown of *wdp* increases the cell surface accumulation of Smo (Fig. 6B). This result suggests that Wdp downregulates Hh signaling either by disrupting Smo translocation to the cell membrane or the stability of Smo on the cell surface.

**Figure 6.**
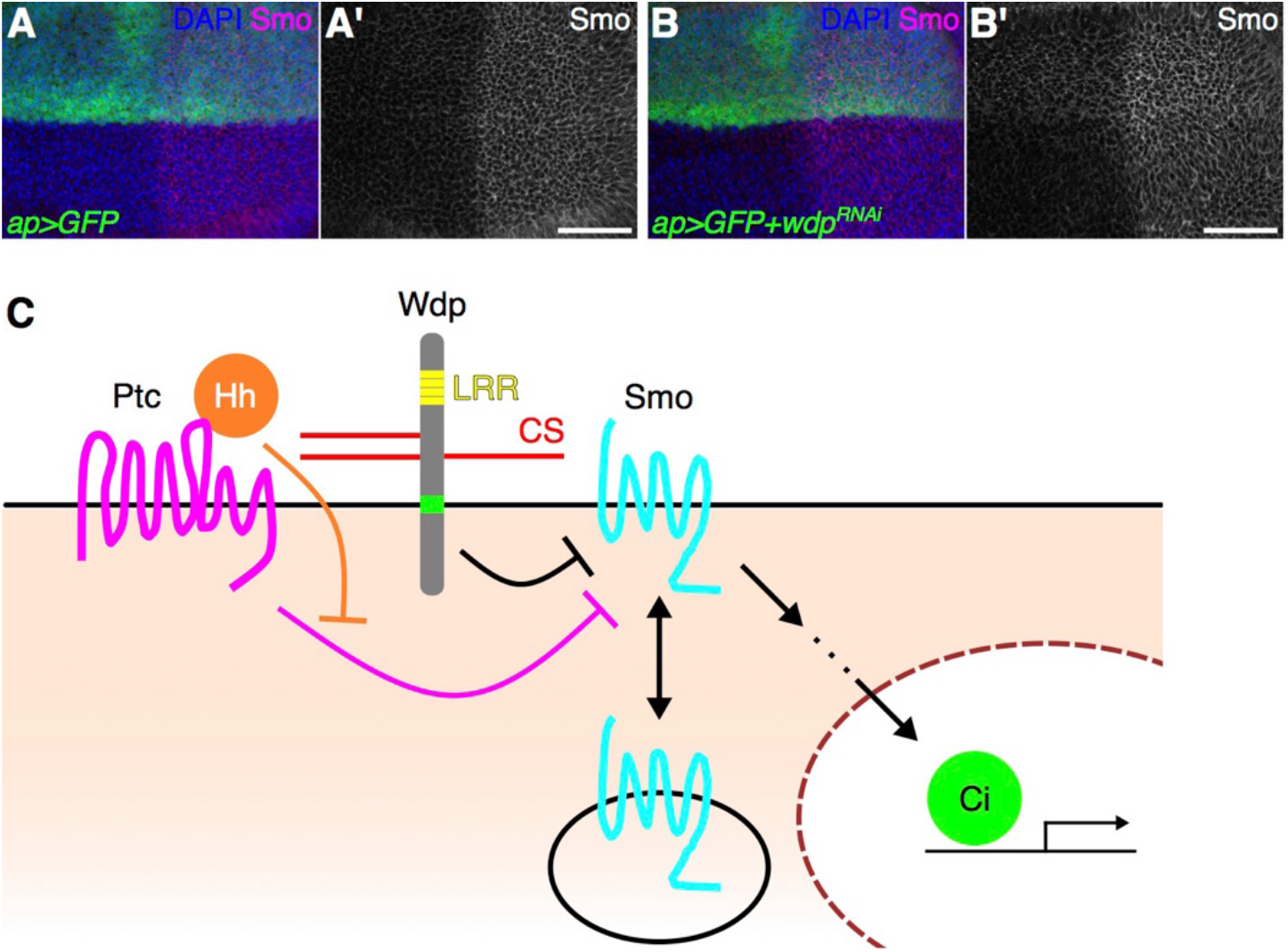
Wdp negatively regulates Smo cell surface accumulation. (**A**, **B**) A control wing disc and a wing disc expressing *UAS-wdp^RNAi^* (TRiP.HMC06302) with *ap-GAL4* were immunostained for the expression of Smo. Nuclei were stained with DAPI. Scale bars: 50 µm. (**C**) A model for how Wdp modulates Hh signaling. Wdp inhibits Smo cell surface accumulation thereby reducing Hh signaling activity.

## Discussion

The molecular mechanism by which HSPG co-receptors regulate growth factor signaling remains a central question in cell biology. Dally, a *Drosophila* HSPG of the glypican type, potentiates Dpp signaling by stabilizing the ligand-receptor complex on the cell surface (Akiyama et al., 2008), suggesting that controlling the rate of receptor-mediated internalization of the signaling complex is the basis for co-receptor activity. However, it is still unknown how HSPGs affect endocytosis and internalization. Since glypicans do not have an intracellular domain, it is likely that these molecules cooperate with other factors (e.g. membrane proteins) to exert co-receptor activity. Thus, it is clear that there are many more unknown factors involved in molecular recognition events on the cell surface. To understand the molecular basis for cell communications, it is critical to identify novel cell surface players.

We found that in the wing disc, Wdp negatively regulates Hh signaling in a CS- and LRR motif-dependent manner. It has also been reported that Wdp negatively regulates JAK–STAT signaling and controls adult midgut homeostasis and regeneration (W. Ren et al., 2015). The authors showed that Wdp interacts with the Dome receptor and promotes its endocytosis and lysosomal degradation. Thus, it is interesting to test if Wdp interacts with Dome via CS chains to modulate JAK–STAT signaling. We observed that Wdp affects cell surface accumulation of Smo, suggesting its role in regulating the stability of Smo protein. Thus, it is possible that Wdp modulates these pathways via a similar mechanism: controlling the internalization of Dome and Smo on the cell membrane.

It is worth noting that both JAK–STAT and Hh signaling, the two pathways negatively controlled by Wdp, are also regulated by HSPGs. Dally-like, a glypican family of HSPGs, positively regulates Hh signaling by interacting with Hh and Ptc (Desbordes & Sanson, 2003; M.-S. Kim, Saunders, Hamaoka, Beachy, & Leahy, 2011; Lum, Yao, et al., 2003a; Williams et al., 2010; Yan et al., 2010). In the developing ovary, Dally upregulates the JAK–STAT pathway (Y. Hayashi et al., 2012). Given the importance of precise dosage control of oncogenic pathways, such as JAK–STAT and Hh signaling, this dual proteoglycan system could play an important role in fine-tuning of the signaling output in order to prevent cancer formation. In vertebrates, HSPGs and CSPGs show opposing effects in neural systems. For example, axon growth is typically promoted by HSPGs but inhibited by CSPGs (Bandtlow & Zimmermann, 2000; Coles et al., 2011; Kantor et al., 2004; Matsumoto, Irie, Inatani, Tessier-Lavigne, & Yamaguchi, 2007; Silver & Miller, 2004; Van Vactor, Wall, & Johnson, 2006). Our findings suggest that such competing effects of HSPGs and CSPGs may be a general mechanism for the precise control of signaling cascades and pattern formation.

In addition to the functions in signaling, Wdp may play other roles. We found that overexpression of *wdp* results in massive apoptosis in the wing disc, independent of Hh signaling inhibition (Fig. S2). Since CSPGs are well known for structural functions, an excess amount of Wdp may affect the epithelial integrity of the wing disc, leading to subsequent apoptosis. Our observation that Wdp is enriched on the basal side of the wing disc and adult midgut cells (Fig. 4B and S3F) suggests that Wdp may interact with components of the basement membrane, which surrounds these organs.

In mice, sulfated CS is necessary for Indian hedgehog (Ihh) signaling in the developing growth plate (Cortes et al., 2009). Ihh and Sonic hedgehog (Shh) bind to CS (Cortes et al., 2009; Whalen, Malinauskas, Gilbert, & Siebold, 2013; F. Zhang, McLellan, Ayala, Leahy, & Linhardt, 2007). Thus, it will be interesting to check if Wdp interacts with Hh via its CS chains.

Previous studies also reported that *wdp* is associated with aggressive behaviors in *Drosophila* species. *wdp* is upregulated in the head of socially isolated male flies, which exhibit more aggressive behaviors than males raised in groups (L. Wang, Dankert, Perona, & Anderson, 2008). Also, *wdp* expression is slightly higher in the brain of *Drosophila prolongata*, which is more aggressive compared to its closely-related species (Kudo et al., 2017). Since CSPGs are important in neuronal patterning (Saied-Santiago & Bülow, 2018), it is interesting to study the molecular mechanisms behind Wdp’s effect on *Drosophila* behavior.

In mammals, there are a class of CSPG molecules with LRR motifs (small leucine-rich proteoglycans, or SLRPs). A number of SLRP members are known as causative genes of human genetic disorders (Bech-Hansen et al., 2000; Pusch et al., 2000; Schaefer & Iozzo, 2008). Although Wdp does not have cysteine-rich regions that are commonly found in mammalian SLRPs, MARRVEL (ver 1.1) (J. Wang et al., 2017) reports that *wdp* is a potential *Drosophila* ortholog of the human *NYX* gene (nyctalopin), a member of SLRPs (DIPOT score 1 (Hu et al., 2011)). Mutations in *NYX* cause X-linked congenital stationary night blindness (Bech-Hansen et al., 2000; Pusch et al., 2000). Further studies on Wdp will provide a novel insight into the function of these disease-related human counterparts.

## Materials and Methods

### Preparation of glycosaminoglycan-glycopeptides and LC-MS/MS analysis

Glycosaminoglycan-glycopeptide samples were prepared from wild-type (Oregon-R) and *ttv* mutant (*ttv^524^*) third-instar larvae as previously described (Noborn et al., 2015; 2018). Briefly, 200-400 third instar larvae (wet weight; 200-400 mg) were lyophilized and homogenized using a motor pestle in 1 ml of ice-cold acetone. After extensive washes with acetone, the insoluble fraction was recovered by centrifugation. After overnight desiccation, the pellet was dissolved in 1.5 ml 1% CHAPS lysis buffer and boiled for 10 min at 96°C. The sample was adjusted to 2 mM MgCl_2_ and incubated with 3 μl Benzonase (MilliporeSigma, Burlington, MA) at 37°C for three hours. After heat-inactivation of Benzonase, the sample was centrifuged and the supernatant was collected in a new tube.

An aliquot of the preparation (1 mg of protein) was further used. The sample was reduced and alkylated in 1 ml 50 mM NH_4_HCO_3_, and trypsinized at 37°C overnight with 20 μg trypsin (Promega, Madison, WI). The digested samples were applied onto DEAE (GE Healthcare, Chicago, IL) columns (600 μl) at 4°C. The columns were washed with three different low-salt washing solutions at 4°C: 50 mM Tris-HCl, 100 mM NaCl, pH 8.0; 50 mM NaAc, 100 mM NaCl, pH 4.0; and 100 mM NaCl. The glycopeptides that were bound to DEAE were eluted stepwise with three buffers with increasing sodium chloride concentrations at 4°C: 4 ml 250 mM NaCl, 400 mM NaCl, 800 mM NaCl, and 3 ml 1500 mM NaCl. Each fraction was desalted using PD10-columns (GE Healthcare).

All fractions were lyophilized and the salt-free samples were then individually treated with 1 mU of chondroitinase ABC (Sigma-Aldrich, St. Louis, MO) for 3 h at 37°C. Prior to MS-analysis, the samples were desalted using a C18 spin column (8 mg resin) according to the manufacturer’s protocol (Thermo Fisher Scientific, Waltham, MA). LC-MS/MS analysis was performed as previously described (Noborn et al., 2015; 2018). In brief, the samples were analyzed on a Q Exactive mass spectrometer coupled to an Easy-nLC 1000 system (Thermo Fisher Scientific). Briefly, glycopeptides (2-μl injection volume) were separated using an analytical column with Reprosil-Pur C18-AQ particles (Dr. Maisch GmbH, Ammerbuch, Germany). The following gradient was run at 300 nl/min; from 7-35 % B-solvent (acetonitrile in 0.2% formic acid) over 75 min, to 100 % B-solvent over 5 min, with a final hold at 100% B-solvent for 10 min. The A-solvent was 0.2% formic acid. Spectra were recorded in positive ion mode and MS scans were performed at 70,000 resolution with a mass range of *m*/*z* 600–1800. The MS/MS analysis was performed in a data-dependent mode, with the top ten most abundant charged precursor ions in each MS scan selected for fragmentation (MS2) by higher energy collision dissociation with normalized collision energy values of 30. The MS2 scans were performed at a resolution of 35,000 (at *m*/*z* 200). The data analyses were performed as previously described (Noborn et al., 2015) with some small adjustments. In brief, the HCD.raw spectra were converted to Mascot .mgf format using Mascot distiller (version 2.3.2.0, Matrix Science, London, UK). The ions were presented as singly protonated in the output Mascot file. Searches were performed using an in-house Mascot server (version 2.3.02) with the enzyme specificity set to *Trypsin*, and then to *Semitrypsin*, allowing for one or two missed cleavages, in subsequent searches on *Drosophila* sequences of the UniprotKB (42, 507, sequences, 2018-06-18). The peptide tolerance was set to 10 parts per million (ppm) and fragment tolerance was set to 0.01 Da. The searches were allowed to include variable modifications at serine residues of the residual hexasaccharide structure [GlcA(-H_2_O)GalNAcGlcAGalGalXyl-O-] with 0 (C_37_H_55_NO_30_, 993.2809 Da), 1 (C_37_H_55_NO_33_S, 1073.2377 Da), or 2 (C_37_H_55_NO_36_S_2_, 1153.1945 Da) sulfate groups attached.

### Fly husbandry and fly strains, and transgenic flies

The following fly strains were used in this study:

Oregon-R, *w^1118^* (Bloomington Drosophila Stock Center [BDSC] #5905), *ttv^524^* (Takei, 2004), *ap-GAL4* (O’Keefe et al., 1998), *hh-GAL4* (Tanimoto et al., 2000), *Bx^MS1096^-GAL4* (BDSC #8860) (Capdevila & Guerrero, 1994), *AB1-GAL4* (BDSC #1824) (Tavsanli et al., 2004), *elav^C155^>mCD8:GFP* (BDSC #5146) (Lin & Goodman, 1994), *UAS-GFP* (BDSC #1521), *UAS-tdTomato* (BDSC #36327 and #36328), *UAS-FLP* (BDSC #4539 and #4540), *UAS-ptc* (BDSC #44614), *nub-GAL4* (BDSC #25754), *FRT42D 2xUbi-GFP*, *UAS-smo:GFP* (BDSC #44624), *UAS-FLAG:smo^Act^* (BDSC #44621), *UAS-wdp^RNAi^* (TRiP.HMC06302, BDSC #66004), *UAS-wdp^RNAi^* (TRiP.HM05118, BDSC #28907), *UAS-smo^RNAi^* (TRiP.HMC03577, BDSC #53348), *hh-lacZ^P^30* (a gift from Gary Struhl) (J. J. Lee et al., 1992), *dpp-lacZ^10638^* (BDSC #12379) (Zecca, Basler, & Struhl, 1995), *vas-Cas9* (BDSC #55821), *esg-GAL4* (DGRC #113886) (S. Hayashi et al., 2002). The *UAS-wdp*, *UAS-wdp^ΔGAG^*, *UAS-Myc:wdp*, *UAS-Myc:wdp^ΔLRR^s*, *UAS-Myc:wdp^ΔICD^*, *wdp^KO.ΔCDS^*, *wdp^KI.HA^*, *wdp^KI.OLLAS^* flies were generated in this study. A full list of genotypes used in this study can be found in Table S1.

For constructing *UAS-wdp*, *wdp* CDS (corresponding to wdp-RA–E in FlyBase) was inserted into the XhoI- and XbaI-digested pJFRC7 vector (a gift from Gerald Rubin; Addgene # 26220) (Pfeiffer et al., 2010) using NEBuilder HiFi DNA Assembly Master Mix (New England Biolabs [NEB], Ipswich, MA, E2621S). Similarly, *wdp^ΔGAG^* (S282A, S334A, and S336A), *Myc:wdp*, *Myc:wdp^ΔLRRs^*, and *Myc:wdp^ΔICD^* were inserted into the pJFRC7 vector. The UAS transgenic flies were generated using *phiC31* integrase-mediated transgenesis at the ZH-86Fb attP (FBti0076525) integration site. Embryonic injection was performed by BestGene Inc (Chino Hills, CA). Primers used in this study will be available upon request.

To generate the *wdp^KO.ΔCDS^* allele, two sgRNAs (pU6-sgRNA-wdp-1 and pU6-sgRNA-wdp-2) were introduced to delete the *wdp* CDS. To construct sgRNA plasmids, 5’ - CTTCGACAGGGCCAACCAGGCGGTC - 3’ and 5’ - AAACGACCGCCTGGTTGGCCCTGTC - 3’ were annealed (pU6-sgRNA-wdp-1); and 5’ - CTTCGAGTGGCCATTGATCACCTGG - 3’ and 5’ - AAACCCAGGTGATCAATGGCCACTC - 3’ (pU6-sgRNA-wdp-2) were annealed and ligated in the BbsI-digested pU6-BbsI-chiRNA plasmid (a gift from Melissa Harrison, Kate O’Connor-Giles, and Jill Wildonger; Addgene #45946) (Gratz et al., 2013). A mixture of 50 ng/µl of pU6-sgRNA-wdp-1 and pU6-sgRNA-wdp-2 was injected into the embryos of the *vas-Cas9* flies, which express Cas9 under the control of the germline *vasa* regulatory elements (Gratz et al., 2014), by BestGene Inc. The *wdp^KO.ΔCDS^* allele was screened by PCR and verified by Sanger sequencing.

To generate the *wdp^KI.HA^* allele, we constructed a donor plasmid, which contained a Gly-Gly-Ser linker, smGFP-HA, and approximately 1-kb homology arms to *wdp* flanking the linker and smGFP-HA, for homology-directed repair. smGFP-HA and the *wdp* homology sequences on either side of the targeted DSB were PCR-amplified from pJFRC201-10XUAS-FRT>STOP>FRT-myr:smGFP-HA (a gift from Gerald Rubin; Addgene plasmid #63166) (Nern et al., 2015) and genomic DNA extracted from the *vas-Cas9* flies, respectively. These fragments were cloned into the pHD-DsRed-attP backbone (a gift from Melissa Harrison, Kate O’Connor-Giles and Jill Wildonger; Addgene #51019) (Gratz et al., 2014) using NEBuilder HiFi DNA Assembly Master Mix (NEB, E2621S). Similarly, we generated a donor plasmid with OLLAS tags amplified from pJFRC210-10XUAS-FRT>STOP>FRT-myr:smGFP-OLLAS (a gift from Gerald Rubin; Addgene plasmid #63170) (Nern et al., 2015). A mixture of 50 ng/µl of pU6-sgRNA-wdp-2 and 125 ng/µl of each donor plasmid was injected into the *vas-Cas9* embryos by BestGene Inc. The *wdp^KI.HA^* and *wdp^KI.OLLAS^* alleles were screened by PCR and verified by Sanger sequencing.

Flies were raised on a standard cornmeal fly medium at 25°C unless otherwise indicated.

### Mosaic analysis

The *wdp^KO.ΔCDS^* homozygous clones were generated by FLP/FRT-mediated mitotic recombination (T. Xu & Rubin, 1993). The FLP expression was induced by *nub-GAL4 UAS-FLP*.

### Immunohistochemstry

Third-instar larval imaginal discs were stained as described previously (Takemura & Adachi-Yamada, 2011) with some modifications. Wing discs were dissected from third-instar wandering larvae in phosphate-buffered saline (PBS, pH 7.4) and subsequently fixed in 3.7% formaldehyde in PBS for 15 min at room temperature. After three 10-min washes with PBST (PBS containing 0.1% (vol/vol) Triton X-100 [Sigma, T8532]), the samples were incubated in primary antibodies overnight at 4°C. After three 10-min washes with PBST, the samples were incubated with Alexa Fluor-conjugated secondary antibodies (1:500, Thermo Fisher Scientific) overnight at 4°C or 2 hours at room temperature. After three 10-min washes with PBST, the samples were stained with 1 µg/ml DAPI (Thermo Fisher Scientific, 62248) and subsequently mounted in VECTASHIELD Antifade Mounting Medium (Vector Laboratories, Burlingame, CA, H-1000). F-actin was stained with Alexa Fluor 568 phalloidin (Thermo Fisher Scientific, A12380). Adult midguts were dissected and immunostained as previously described (Takemura & Nakato, 2017). Images were acquired on a LSM710 confocal microscope (Carl Zeiss, Oberkochen, Germany). For quantification of Ptc staining, images were acquired with the same condition, and fluorescence intensity was measured in a set area with Fiji (Schindelin et al., 2012).

### Antibodies

The primary antibodies used were as follows: mouse anti-Ptc Apa 1 (1:20, Developmental Studies Hybridoma Bank [DSHB], Iowa City, IA, deposited by Isabel Guerrero) (Capdevila et al., 1994), rat anti-Ci 2A1 (1:20, DSHB, deposited by Robert Holmgren) (Motzny & Holmgren, 1995), chicken anti-βGalactosidase (1:2000, Abcam), mouse anti-En 4D9 (1:20, DSHB, deposited by Corey Goodman) (Riggleman, Schedl, & Wieschaus, 1990), rabbit anti-pH3 (1:1000, Millipore, 06-570), rat anti-HA 3F10 (1:200, Roche, 11867423001), rabbit anti-HA C29F4 (1:1000, Cell Signaling, 3724), mouse anti-Smo 20C6 (1:50, DSHB, deposited by Philip Beachy) (Lum, Zhang, et al., 2003b), rabbit anti-pSmad3 (1:1000, Epitomics, 1880-1) (Smith, Machamer, Kim, Hays, & Marques, 2012), rabbit anti-Salm (1:30, a gift from Scott Selleck), mouse anti-Dll 1:500 (1:500, a gift from Dianne Duncan) (D. M. Duncan, Burgess, & Duncan, 1998), guinea pig anti-Sens (1:1000, a gift from Hugo Bellen) (Nolo, Abbott, & Bellen, 2000), rabbit anti-cleaved Caspase-3 (1:200, Cell Signaling, 9661), rat anti-OLLAS L2 (1:500, Novus Biologicals, NBP1-06713), mouse anti-Arm N2 7A1 (1:50, DSHB, deposited by Eric Wieschaus) (Riggleman et al., 1990), mouse anti-Pros MR1A (1:50, DSHB, deposited by C.Q. Doe) (Campbell et al., 1994), and mouse anti-Fas3 7G10 (1:50, DSHB, deposited by Corey Goodman) (Patel, Snow, & Goodman, 1987). Alexa488, Alexa548, Alexa564 and Alexa633-conjugated secondary antibodies (Thermo Fisher Scientific) were used at a dilution of 1:500.

### Adult wing preparation

The left wings from female flies were dissected and mounted on slides using Canada balsam (Benz Microscope, BB0020) as previously described (Takemura & Adachi-Yamada, 2011). Images were taken using a ZEISS Stemi SV 11 microscope equipped with a Jenoptic ProgRes C3 digital camera.

## Acknowledgements

We thank Melissa Harrison, Kate O’Connor-Giles, Jill Wildonger, Gary Struhl, Scott Selleck, Hugo Bellen, the Bloomington Drosophila Stock Center (NIH P40OD018537), the Transgenic RNAi Project at Harvard Medical School [NIH/NIGMS R01-GM08947], and the Drosophila Genomics Resource Center (NIH 2P40OD010949) for sharing fly strains and plasmids. We also thank the Proteomics Core Facility at the Sahlgrenska Academy, University of Gothenburg, Sweden for running all the MS analyses. This work was supported by the National Institutes of Health (R01 GM115099) to H.N. M.T. held postdoctoral fellowships from the Japan Society for the Promotion of Science and the Uehara Memorial Foundation.

---

**Figure S1.**
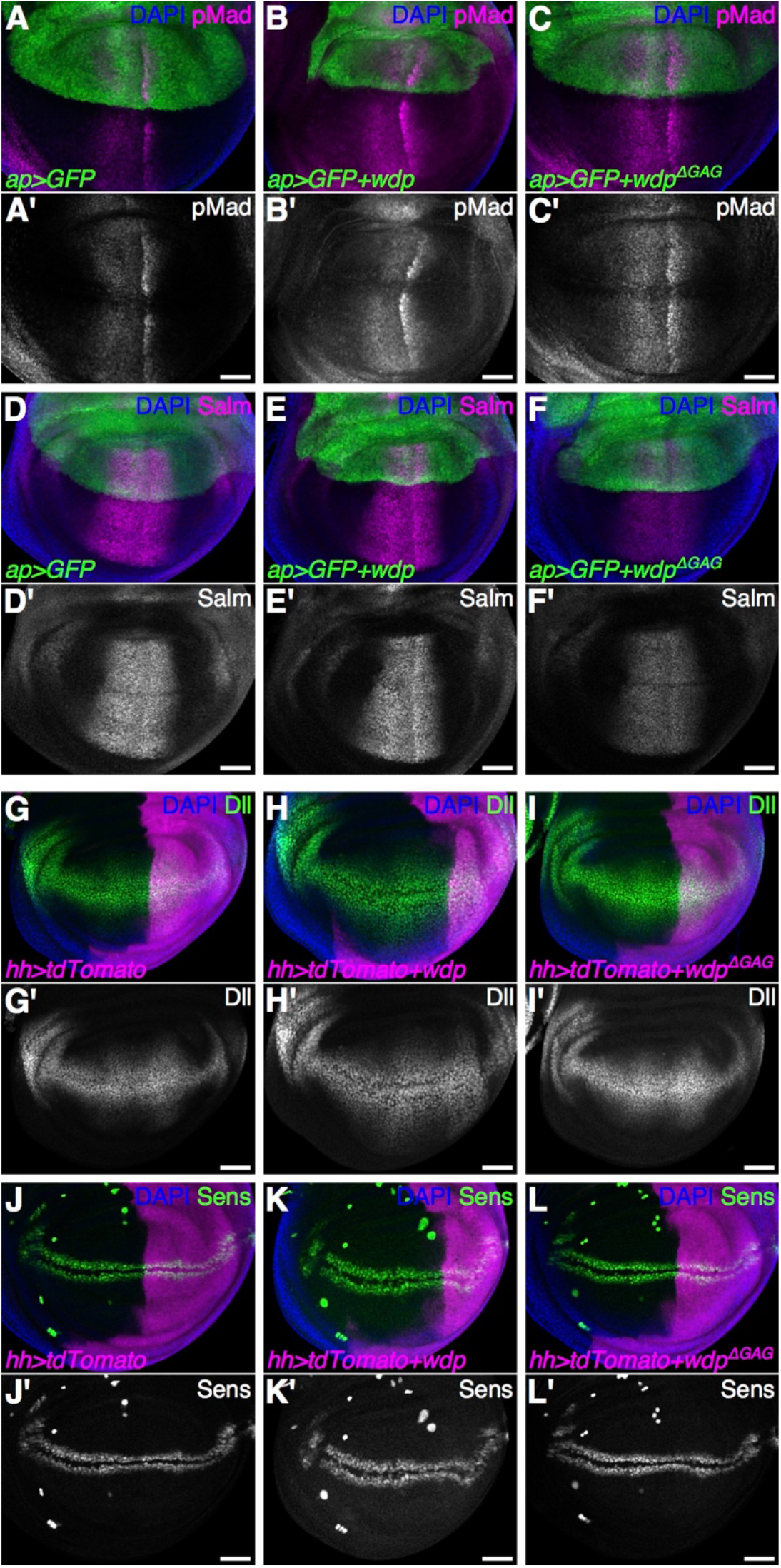
Effect of *wdp* overexpression on Dpp and Wg signaling. (**A**–**F**) Control wing discs (A and D) and wing discs expressing *UAS-wdp* (B and E) or *UAS-wdp^ΔGAG^* (C and F) with *ap-GAL4* were immunostained for pMad (A–C) and Salm (D–F) as read-outs of Dpp signaling. (**G**–**L**) Control wing discs (G and J) and wing discs expressing *UAS-wdp* (H and K) or *UAS-wdp^ΔGAG^* (I and L) with *hh-GAL4* were immunostained for Dll (G–I) and Sens (J–L) as read-outs of Wg signaling (Nolo et al., 2000; Zecca, Basler, & Struhl, 1996). Nuclei were stained with DAPI. Scale bars: 50 µm.

**Figure S2.**
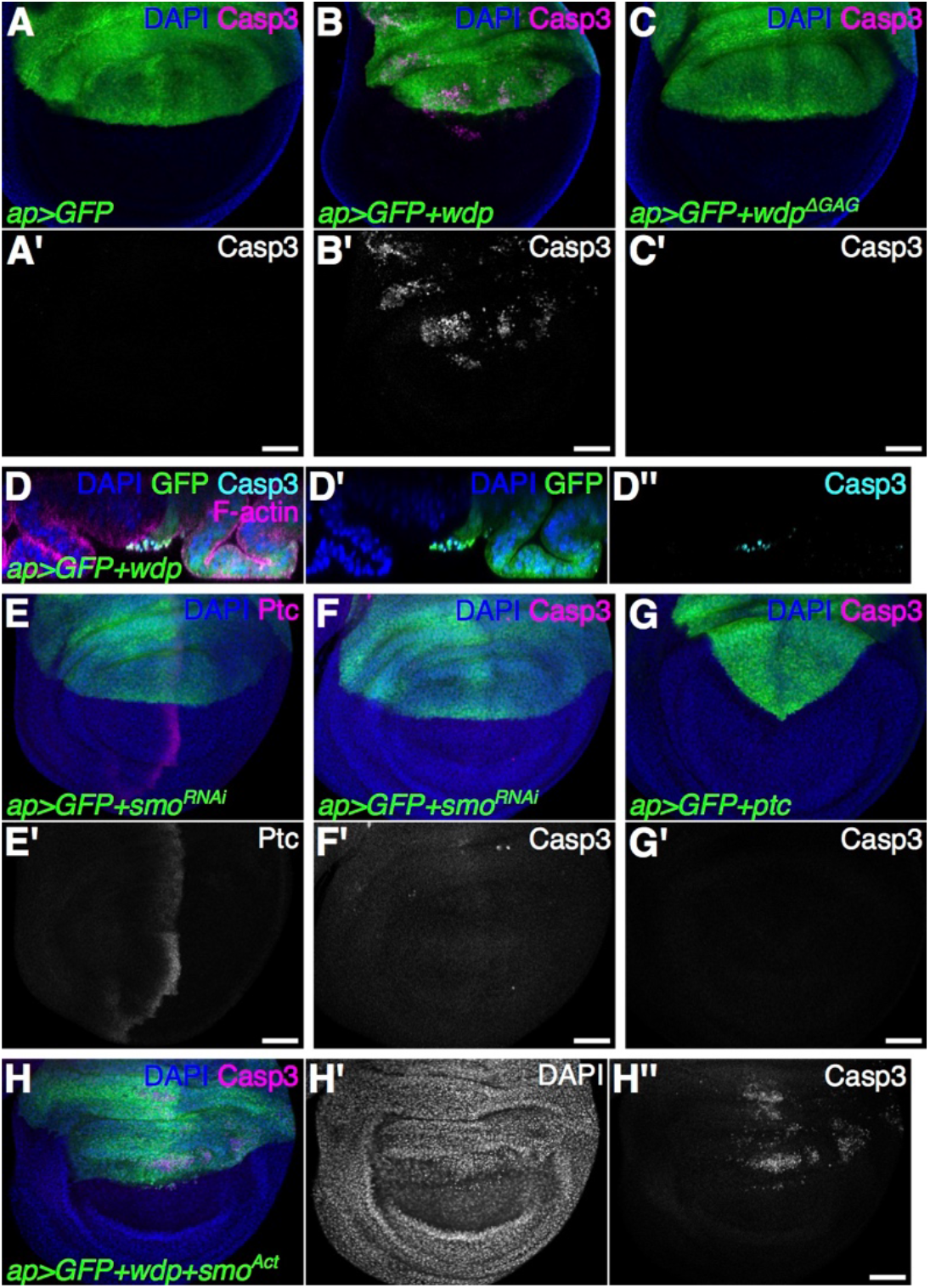
Overexpression of *wdp* in the wing disc induces apoptosis. (**A–C**) A control wing disc (A) and wing discs expressing *UAS-wdp* (B) or *UAS-wdp^ΔGAG^* (C) with *ap-GAL4* were immunostained for cleaved Caspase-3 as a marker of apoptotic cells. (**D**) An optical cross section of a wing disc expressing *UAS-wdp* with *ap-GAL4*. Signals for cleaved Caspase-3 (cyan) and F-actin (magenta) are shown. (**E**) A wing disc expressing *UAS-smo^RNAi^* (TRiP.HMC03577) with *ap-GAL4* was immunostained for Ptc. (**F** and **G**) Wing discs expressing *UAS-smo^RNAi^* (F) or *UAS-ptc* (G) with *ap-GAL4* were immunostained for cleaved Caspase-3. (**H**) A wing disc coexpressing *UAS-wdp* and *UAS-FLAG:smoAct* was immunostained for cleaved Caspase-3. Nuclei were stained with DAPI. Scale bars: 50 µm.

**Figure S3.**
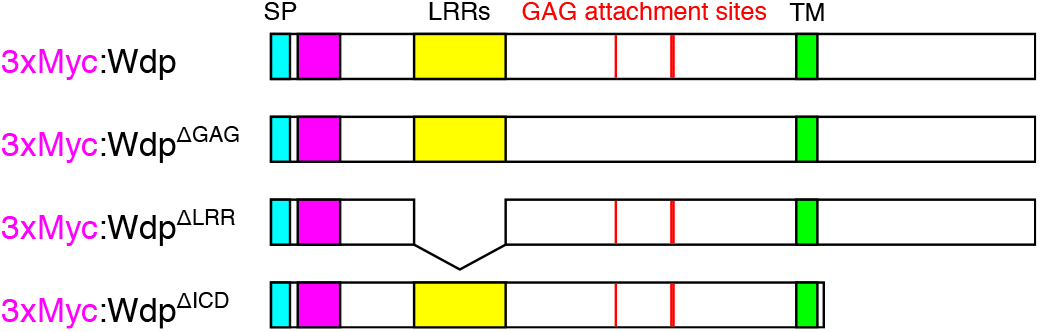
A schematic of 3xMyc-tagged wild-type and mutant Wdp constructs. The magenta box denotes 3 copies of a Myc tag. The cyan box indicates the signal peptide (SP). The yellow box shows the LRR motifs. The red line indicates the CS attachment sites. The green box shows the transmembrane domain (TM).

**Figure S4.**
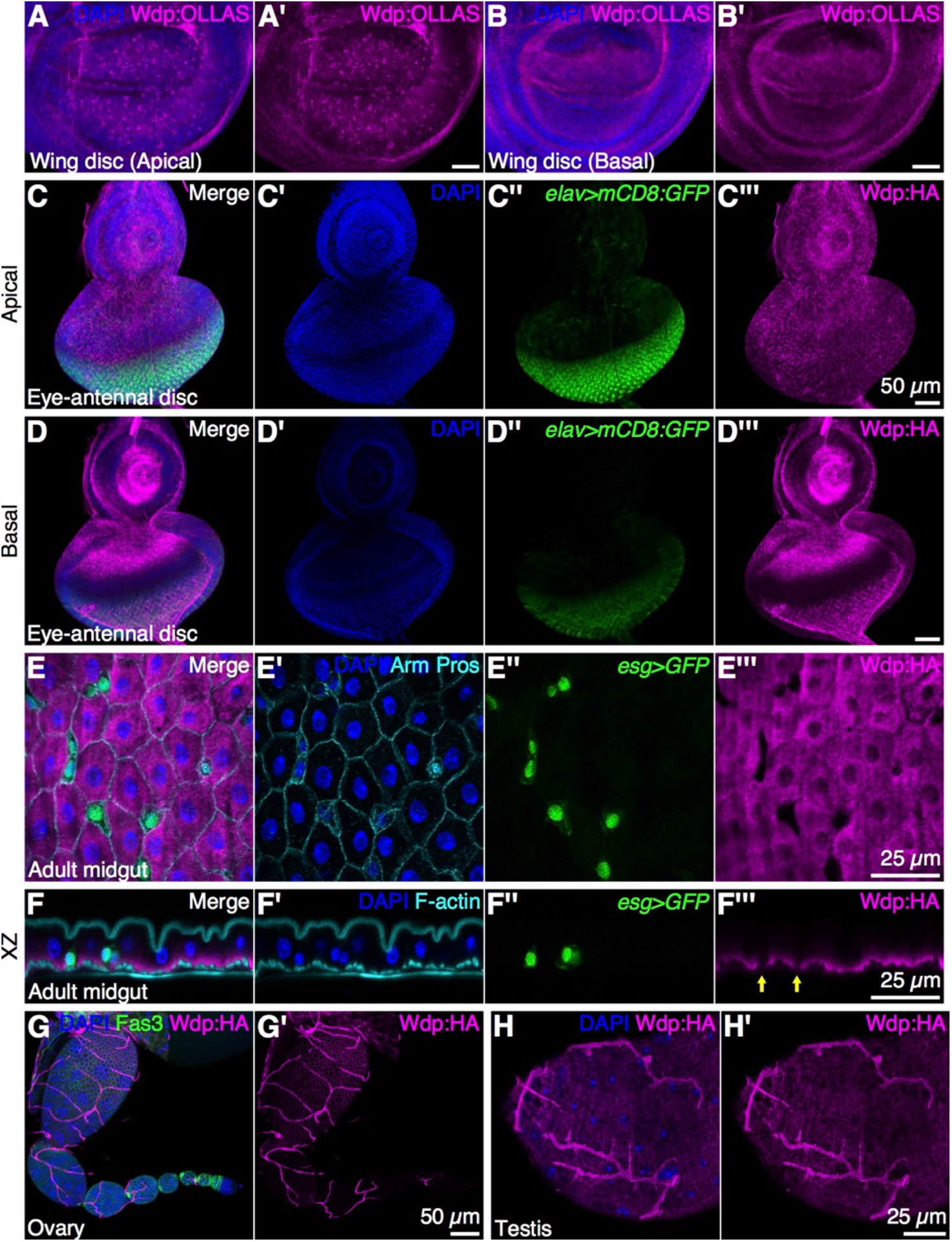
Expression patterns of Wdp in the eye disc, adult posterior midgut, ovary and testis. (**A**,**B**) Apical (A) and basal (B) sections of a wing disc homozygous for *wdp^KI.OLLAS^* was stained with anti-OLLAS antibody. (**C**–**H**) An eye disc, adult midgut, ovary, and testis with one or two copies of *wdp^KI.HA^* were immunostained with anti-HA antibody (magenta). (C, D) Apical (C) and basal (D) sections of an eye disc. Neurons are marked by *elav-GAL4 UAS-mCD8:GFP* (*elav>mCD8:GFP*, green). (E) An adult midgut showing ISCs/enteroblasts (*escargot-GAL4 UAS-GFP* [*esg>GFP*]), enteroendocrine cells (nuclear Prospero [Pros]), cell membrane (Armadillo [Arm]), and Wdp:HA. (F) An optical cross section of an adult midgut showing ISCs/enteroblasts (*esg>GFP*) and F-actin (stained by phalloidin-Alexa568). (G) Wdp:HA expression is detected in the egg chambers (stage 6 and later) in the ovary. Wdp:HA is also detectable in the tracheal system. (H) Wdp:HA expression is detected in the sheath cells and the tracheal system in the testis. Nuclei were stained with DAPI. Scale bars: 50 µm (A, B, C, D, and G), 25 µm (E, F, and H).

**Table S1.**
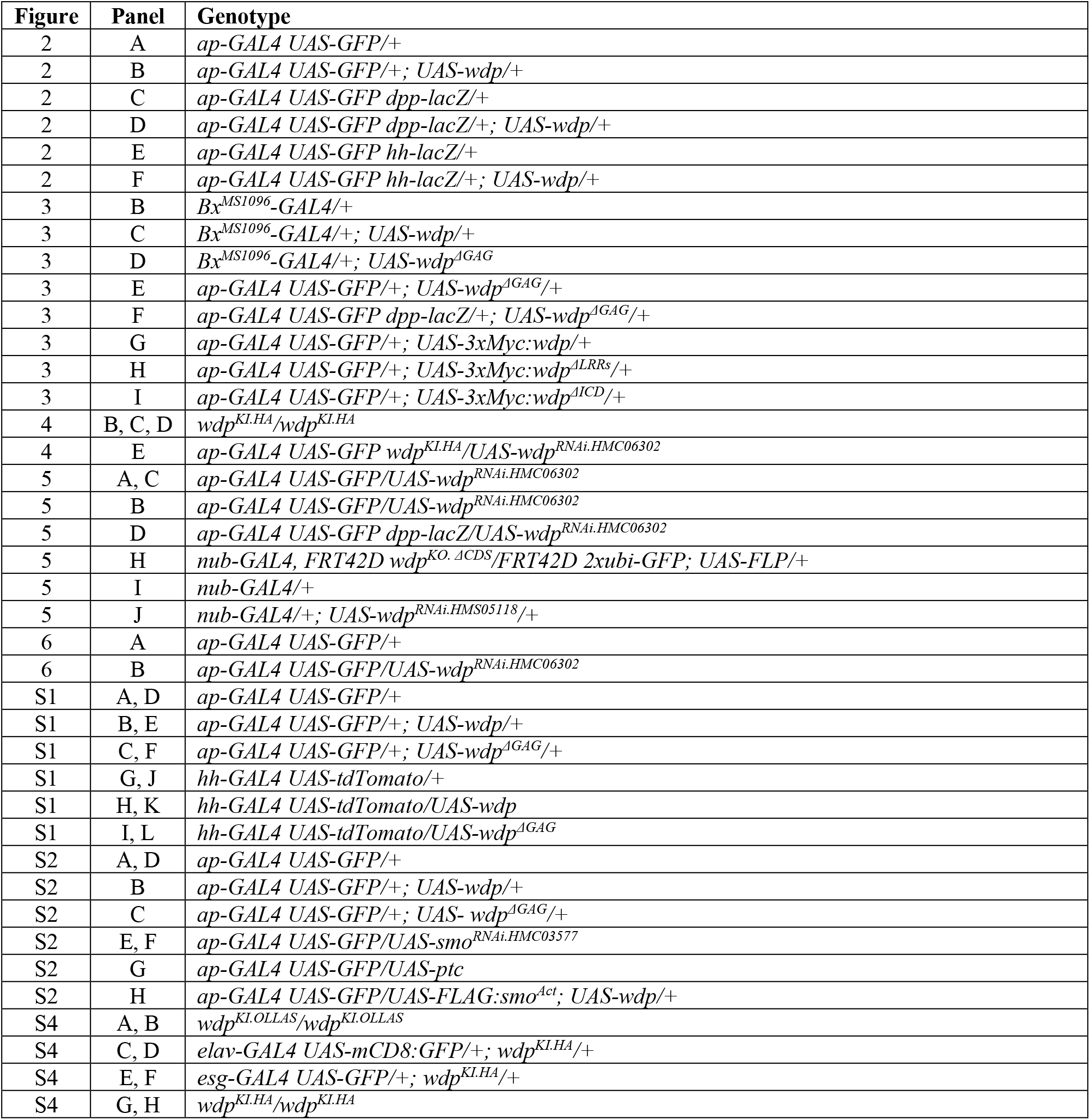
Genotypes of *Drosophila* strains used in each figure.

